# COVID-19 dominant D614G mutation in the SARS-CoV-2 spike protein desensitizes its temperature-dependent denaturation

**DOI:** 10.1101/2021.03.28.437426

**Authors:** Tzu-Jing Yang, Pei-Yu Yu, Yuan-Chih Chang, Chu-Wei Kuo, Kay-Hooi Khoo, Shang-Te Danny Hsu

**Affiliations:** Institute of Biological Chemistry, Academia Sinica, Taipei 11529, Taiwan; Institute of Biochemical Sciences, National Taiwan University, Taipei 10617, Taiwan; Academia Sinica Cryo-EM Center, Academia Sinica, Taipei 11529, Taiwan

## Abstract

The D614G mutation in the spike protein of SARS-CoV-2 alters the fitness of the virus, making it the dominant form in the COVID-19 pandemic. Here we demonstrated by cryo-electron microscopy that the D614G mutation does not significantly perturb the structure of the spike protein, but multiple receptor binding domains are in an upward conformation poised for host receptor binding. The impact of the mutation lies in its ability to eliminate the unusual cold-induced unfolding characteristics, and to significantly increase the thermal stability under physiological pH. Our findings shed light on how the D614G mutation enhances the infectivity of SARS-CoV-2 through a stabilizing mutation, and suggest an approach for better design of spike-protein based conjugates for vaccine development.

## Main text

The COVID-19 (coronavirus disease 2019) pandemic is caused by the infection of SARS-CoV-2 (severe acute respiratory syndrome coronavirus 2).^1^ Early bioinformatic analysis of the reported genome sequences of SARS-CoV-2 revealed the emergence of a prominent pairwise linkage disequilibrium between three nucleotide changes since mid-February, 2020, namely nt3037 (C > T), nt14408 (C > T), and nt23403 (A > G). The last mutation corresponds to a missense D614G mutation in the spike (S) protein.^2^ The G clade, which harbors the aforementioned three nucleotide changes and the nt241 (C > T) mutation, relative to the original Wuhan form (hereafter wild type, WT), became the dominant form of the COVID-19 pandemic since the summer of 2020.^3^ The D614G variant is found to exhibit enhanced infectivity of in both cell culture and animal models.^4^

The SARS-CoV-2 S protein (hereafter S protein) is responsible for host recognition and viral entry. The S protein binds to the receptor, angiotensin converting enzyme 2 (ACE2), through an upward open RBD conformation (RBD-up); a downward closed RBD conformation (RBD-down) sequesters its receptor binding motif (RBM) from receptor binding, rendering such a conformation inactive.^5-6^ Several studies on COVID-19 convalescent sera have identified multiple antibodies that competitively bind to the RBM, and in doing so prevent host receptor ACE2 binding, thereby achieving neutralizing activities.^7-11^ Despite the large size, the S protein is marginally stable over a narrow range of temperatures. Incubation of the recombinant SARS-CoV-2 S protein at 4 °C leads to significant unfolding within 24 hours. The morphological change of cold denaturation resembles that after a brief heat shock at 50-60 °C.^12-13^ This instability complicates the use the S protein as an antigen for vaccine development. Thus, efforts have been made towards engineering more thermal stable SARS-CoV-2 S protein variants.^12, 14-15^ One study provides a simple solution to revitalize the cold-denatured WT S protein (hereafter S-D614) by incubation at 37 °C.^13^ There is good evidence to suggest that the S protein of the D614G variant (hereafter S-D614G) is more thermally stable than S-D614.^16^ Several cryo-electron microscopy (cryo-EM) studies indicate increased conformational heterogeneity and more RBD-up conformations of S-D614G RBD dynamics, which may be implicated in the altered ACE2 binding kinetics.^17-20^

Here we characterized the structure and dynamics of S-D614G by cryo-EM aided by three dimensional variability analysis (3DVA).^21^ We identified five distinct but equally populated clusters of conformations of S-D614G with varying degrees of RBD-up populations (Fig. 1, Fig. S1 and Table S1). Three of the five clusters were in one RBD-up states and the remaining two clusters were in two RBD-up states. Contrary to S-D614, we did not observe any all RBD-down conformation in S-D614G, in line with the previous study of a similar construct without the transmembrane domain^18^, but differs from the full-length S-D614G that exhibits significant amount of closed, all RBD-down conformation.^19-20^ A common feature of reported S-D614G structures, however, is the presence of two RBD-up conformation that is absent in S-D614.

**Fig. 1.**
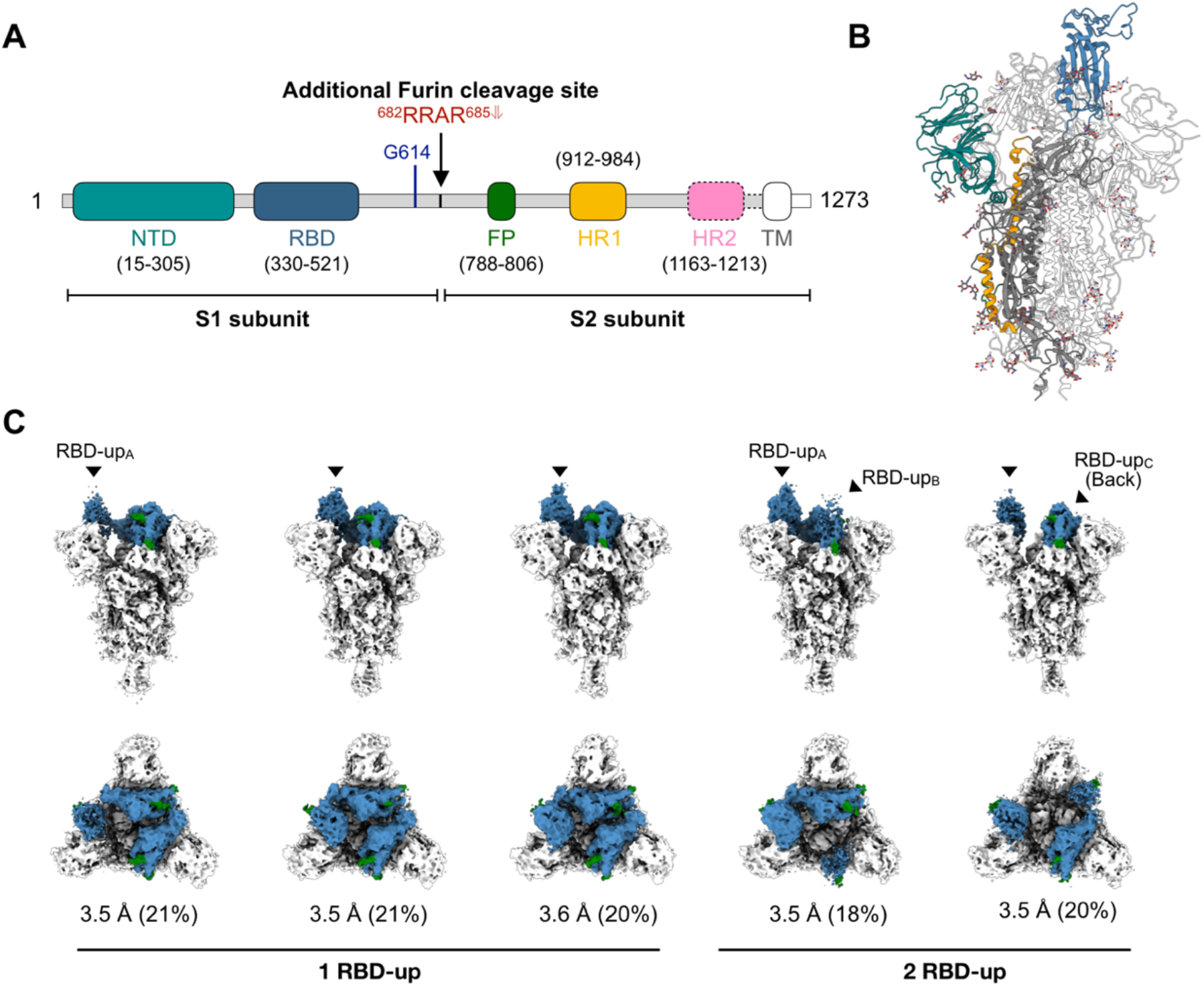
Cryo-EM analysis of S-D614G. (A) Schematic domain architecture of S-D614G. NTD, N-terminal domain. RBD, receptor-binding domain. FP, fusion peptide. HR1/HR2, heptad repeat 1/2. TM, transmembrane domain. The regions not resolved in the cryo-EM map are highlighted with dash line. (B) Cartoon representative of the atomic model of the trimeric S-D614G with the domains colored in accordance with (A). (C) Structural heterogeneity of S-D614G. Orthogonal views – side views and top views are shown on the top and bottom panels, respectively – of the five distinct clusters of S-D614G derived from 3DVA. The RBDs are colored in blue, and the two N-glycans on the RBD, N331 and N343, are colored in green. The nominal resolution of the cryo-EM map and the relative population in percentages are shown below each cluster.

Yurkovetskiy *et al*. proposed that the D614G mutation may disrupt the inter-protomer hydrogen bond between D614 of one protomer and T859 of the other protomer thereby leading to increase dynamics of the S protein and the shift of equilibrium between the up-RBD and down-RBD.^17^ However, our cryo-EM structure of S-D614G showed no appreciable structural difference in the proximity of the mutation site with respect to some of the extremities of the reported S-D614 structures, including the first reported structures of S-D614^5-6^, the acid-stabilized form of S-D614^16^, the disulfide- and proline-stabilized HexPro-S variant^15^, and the furin-cleaved S-D614 that adopts a more open conformation (Fig. S2-3).^22^ Meanwhile, Benton *et al*. proposed that the loss of a key salt bridge formed by D614 of one protomer and K584 or the other protomer due to the D614G mutation leads to local disorder thereby increasing the dynamics of S-D614G and more populated RBD-up conformations^18^, a finding that could be confirmed by our cryo-EM structures (Fig. S4). Nevertheless, the molecular basis of how the D614G mutation could allosterically change the conformation and dynamics of the RBDs remains to be established.

Considering that the D614G mutation is adjacent to the N-glycosylation site of N616, we sought to examine how the mutation may impact on the glycosylation of N616. Using glycopeptide analysis by mass spectrometry (MS), we observed generally consistent glycoforms in both S-D614 and S-D614G as previously reported.^23^ Nevertheless, we observed a significant enrichment of the high-mannose type glycosylation with five mannose (Man) moieties, Man_5_GlcNAc_2_ (M5), and more heterogenous complex type glycoforms on N616 for S-D614G relative to S-D614 (Fig. S5). This finding suggests that the N-glycosylation site exhibits different conformational dynamics during the biogenesis that could alter the interplay between the S protein and the glycosylation machinery.^24^

Having established that the structure of S-D614G is not significantly perturbed by the D614G mutation under native conditions, we sought to investigate the impact of the D614G mutation on the thermal stability over a range of experimental conditions. Knowing that the S protein is susceptible to cold denaturation, all experiments were carried out using freshly prepared S-D614 and S-D614G, which were secreted into the culture media at 37 °C, and purified at room temperature (Fig. S6). We carried out negative stain electron microscopy (NSEM) analyses on S- D614 and S-D614G on day 0, and on day 6 after continuous incubation at 37 °C and 4 °C. In line with the previous finding^13^, S-D614 remained stable at 37 °C for six days without discernible morphological change, while significant unfolding was observed after incubation at 4 °C for six days (Fig. 2A).

**Fig. 2.**
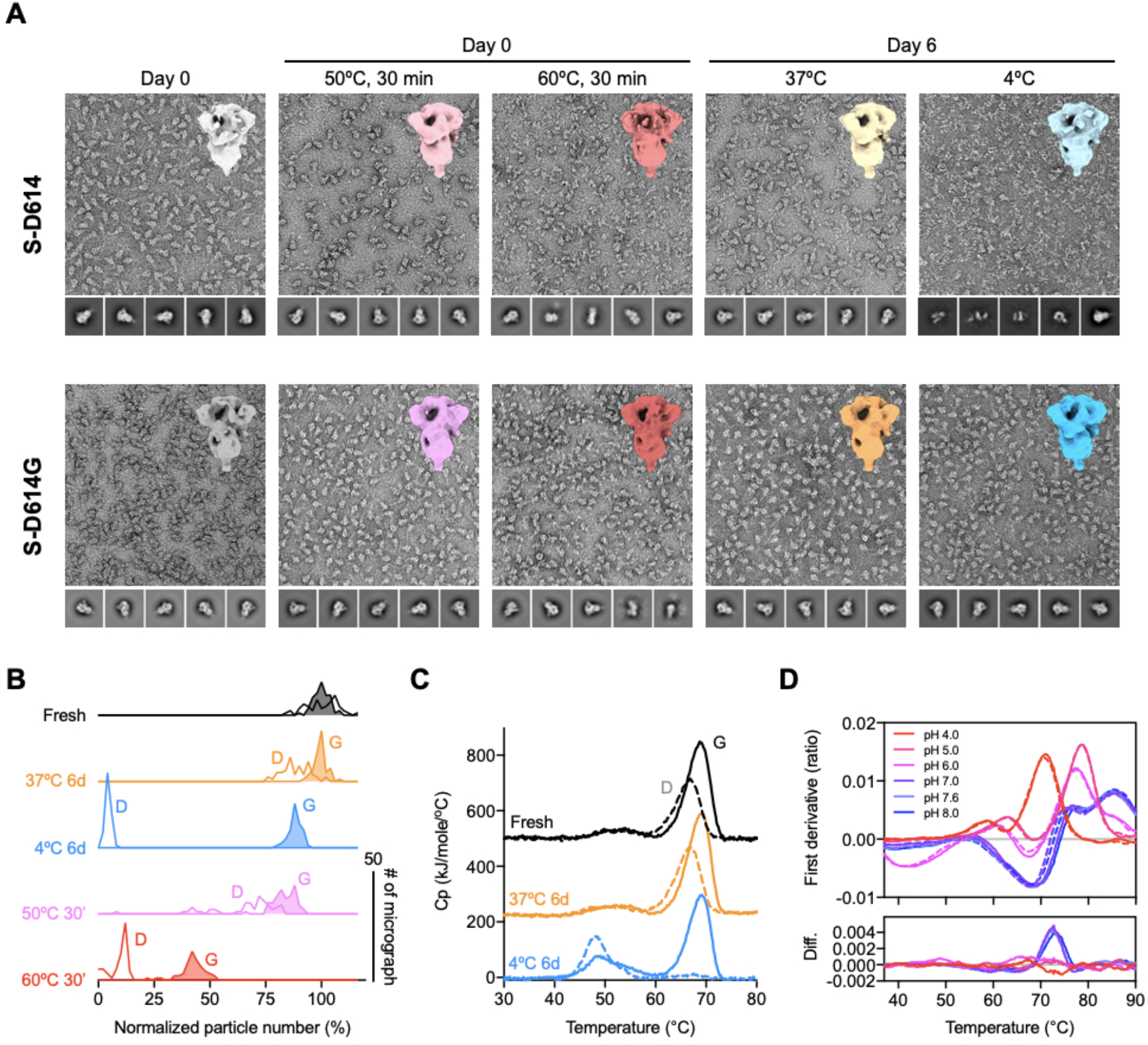
S-D614G is highly stable over a broad range of temperatures. (A) Representative NSEM micrographs of S-D614 (top panels) and S-D614G (bottom panels) after different temperature treatments. Selected 2D classes of picked particle images are shown below each panel. Significant unfolding of S-D614 was observed after heat shocks and after six days of incubation at 4 °C. In contrast, S-D614G exhibited visible unfolding only by heat shock at 60 °C for 30 min. 3D maps of the individual samples derived from the native-like particle images are shown on the upper right corner of each panel. (B) Histograms of the relative amounts of native-like particle images with respect to the fresh samples. The Y-axis correspond to the number of micrographs (n = 50–60). The open and filled curves correspond to S-D614 and S-D614G, respectively. (C) DSC profiles S-D614 (dashed lines) and S-D614G (solid lines) at Day 0 (fresh; black), 37°C for six days (orange) and 4 °C for six days (light blue). (D) DSF profiles of S-D614 (dashed lines) and S- D614G (solid lines) as a function of pH values. The difference between S-D614G and S-D614 (Diff.) is derived by subtracting the values of S-D614 by those of S-D614G.

Quantitative analysis showed that the six-day incubation at 37 °C and 4 °C resulted in 12±6 % and 96±1 % loss of native particles, respectively (Fig. 2A and 2B, Table S2). Unexpectedly, S- D614G did not show significant unfolding after the same cold treatment: the six-day incubation at 37 °C and 4 °C resulted in 1±3 % and 12±3 % loss of native particles, respectively (Fig. 2A and 2B). Remarkably, after two months of incubation at 37 °C, S-D614G remained mostly intact according to NSEM analysis (Fig. S7). When subjected to heat-induced denaturation of both proteins at 50 °C and 60 °C, S-D614 exhibited significant unfolding at 30 min of heat shock at 50 °C and 60°C, resulting in 34±15 % and 91±4 % loss of native particles, respectively. In contrast, the same heat shock treatment resulted in 17±5 % (50 °C) and 58±6 % (60 °C) loss of native particles for S-D614G. Furthermore, S-D614 exhibited more variations in the number of native particles per micrograph, particular in the case of heat treatment at 50 °C for 30 min (Fig. 2B). Note, however, that despite the significant reduction in the number of native-like particles after cold and/or heat treatments, the resulting 3D EM maps of S-D614 and S-D614G were very similar at the resolution of approximately 10 Å (insets in Fig. 2A), suggesting that the unfolding processes of the S protein variants follows an apparent two-state model resulting in different levels of native populations after the treatments. Indeed, further analysis with more incubation temperatures for S- D614G yielded a two-state-like melting curve with an apparent T_m_ of 59 °C (Fig. S8).

We further analyzed the thermal unfolding of S-D614 and S-D614G at pH 7.6 by differential scanning calorimetry (DSC). Both S-D614 and S-D614G exhibited two transition peaks, and shared a similar melting temperature (T_m_) for the first transition at *ca*. 52 °C. However, S-D614G exhibited a higher T_m_ for the second transition (T_m_ = 68.8 °C) compared to that of S-D614 (T_m_ = 66.9 °C). Moreover, the total enthalpy of unfolding ΔH of S-D614G was approximately 40 % higher than that for S-D614 (Fig. 2C and Table S3). The bimodal distribution of the DSC profile of S-D614G has been reported previously^13, 16^, and that the second transition was missing in some cases; this may be attributed to prolonged sample storage at lower temperature.^16^ Indeed, the second transition peak was lost for S-D614 after incubation at 4 °C for six days, while that of S- D614G remained largely unaffected; in line with NSEM results, long-term incubation at 37 °C did not significantly affect both variants (Fig. 2C). While keeping purified proteins at 4 °C for long-term storage is generally considered to be more favorable than storage at 37 °C, this is not applicable to S-D614. In contrast, S-D614G is not sensitive to cold-denaturation, and is more robust during transient heat shock (Table S3). The stabilization effect of the D614G mutation is most pronounced under neutral to slightly alkaline conditions as demonstrated by label-free differential scanning fluorimeter (DSF) analyses of S-D614 and S-D614G over a range of pH values (Fig. 2D). While both variants exhibited multiple inflection temperatures (T_i_) between pH 4 and 8 as has been reported previously^16^, their profiles were indistinguishable at pH 6 and below. For pH 7.0 and above, only the transition between 70 and 75 °C showed a clear difference between the two variants, suggesting that the stabilizing effect of D614G is likely contributing to the integrity of the S protein before being endocytosed into the host cell.

In summary, we determined the cryo-EM structure of S-D614G, and revealed the high degree of conformational heterogeneity mostly confined within the RBDs (Fig. 1C). Contrary to the previous report by Yurkovetskiy *et al*., we did not observe appreciable conformational rearrangement around the inter-protomer interaction site between D614 and T859^17^, but we did observe significant disorder of the K854 loop as reported by Benton *et al*. (Fig. S4).^18^ Furthermore, we observed small but significant increase of the high-mannose type glycans (M5) for N616. The higher abundance of the M5 N-glycans is indicative of a more occluded space around the N-glycosylation sites, which limits the subsequent processing to generate complex type glycans.^24^ The most significant finding is the markedly increased resistance of S-D614G to withstand cold- and heat-induced unfolding (Fig. 2). Although the loss of the salt bridge between D614 and K584 is enthalpically unfavorable, it is compensated by other enthalpic gains evidenced by the significantly higher total enthalpy of unfolding ΔH of S-D614G compared to that of S-D614 (Fig. 2C). We argue that the increased stability of S-D614G could be further attributed to the increased configurational entropy manifested in the higher conformational heterogeneity of the RBDs, which contributes to the free energy gain of the system.

The significance of our findings is three-fold. First, the increased folding stability may help explain the gain in fitness of the G clade SARS-CoV-2, which relies on the surface S protein to recognize host receptor molecules, namely ACE2. The elimination of the cold sensitivity of the S protein will undoubtedly increase the robustness of the infection machinery over a range of environment conditions. Second, our DSC analysis of freshly prepared S protein variants hint at the possibility that some of the previously reported cryo-EM and biophysical studies of the SARS-CoV-2 S protein may have been affected by its sensitivity to cold treatments leading to the unfolding and the loss of structural elements that give rise to the thermal transition peaks at around 60–70 °C (Fig. 2C). More importantly, the partial unfolding of the S protein is expected to significantly alter the accessibility of the RBD to which ACE2 and neutralizing antibodies bind and hence impact interpretations of binding assays. Third, the ability of S-D614G to withstand long-term storage at 4 °C without unfolding the prefusion state provides a solution to better vaccine designs and formulation without the need to introduce a large number of mutations and disulfide bonds.^12, 14-15^

## ASSOCIATED CONTENT

Experimental materials and methods, including molecular cloning, protein expression and purification, NSEM and cryo-EM, DSF and DSC, are described in Supporting Information. The atomic coordinates of the S-D614G are deposited in the Protein Data Bank (PDB) under the accession codes 7EAZ, 7EB0, 7EB3, 7EB4 and 7EB5. The corresponding cryo-EM maps are deposited in the Electron Microscopy Data Bank (EMDB) under accession codes EMD-31047, 31048, 31050, 31051 and 31052.

## Supporting Information

The supporting information includes:

Figures S1-S8, Table S1-S3 and references.

## AUTHOR INFORMATION

### Author Contributions

S.-T.D. H. conceived the experiment. T.J.Y. and P.Y.Y. prepared the samples. T.J.Y., P.Y.Y. C.W.K, and Y.C.C. collected the data. T.J.Y., P.Y.Y. C.W.K., K.H.K. and S.-T.D.H. analyzed the data. T.J.Y. and S.-T.D.H. wrote the manuscript with inputs from all co-authors. S.-T.D.H. obtained funding.

### Funding Sources

This work was supported by Academia Sinica intramural fund to S.-T.D.H and K.H.K, an Academia Sinica Career Development Award, Academia Sinica to S.-T.D.H (AS-CDA-109-L08), and the Ministry of Science and Technology (MOST), Taiwan (MOST 109-3114-Y-001-001 to S.-T.D.H.).

## ACKNOWLEDGMENT

We thank the Academia Sinica Biophysics Core Facility (Grant AS-CFII108-111), Academia Sinica Common Mass Spectrometry Facilities (Grant AS-CFII-108-107), and Academia Sinica Cryo-EM Center (Grant AS-CFII-108-110) for data collection, all of which are funded by the Academia Sinica Core Facility and Innovative Instrument Project. We also thank the mammalian cell culture facility of Institute of Biological Chemistry, Academia Sinica, for supporting the protein production, and the Academia Sinica Grid Computing for cryo-EM data processing.

## Notes

### Competing Interest Statement

The authors have declared no competing interest.

