## Supplementary Materials for "COVID-19 dominant D614G mutation in the SARS-CoV-2 spike protein desensitizes its temperature-dependent denaturation"

**This PDF file includes:**

Materials and Methods
Figs. S1 to S8
Tables S1 to S3

### Materials and Methods

#### Expression and purification of SARS-CoV-2 S and its variant, S-D614G

The codon-optimized nucleotide sequence of full-length SARS-CoV-2 S protein was a kind gift of Dr. Che Alex Ma (Genomics Research Center, Academia Sinica). The DNA sequence corresponding the residues 1-1208 of the S protein was subcloned into the mammalian expression vector pcDNA3.4-TOPO (Invitrogen). Additionally, a tandem proline mutation (2P, $^{986}$KV$^{987}$ → $^{986}$PP$^{987}$) and changes at the furin cleavage site (fm, $^{682}$RRAR$^{685}$ → $^{682}$GSAG$^{685}$) were introduced for stabilization[1], which corresponds to S-D614. The D614G mutation was subsequently introduced to generate S-D614G. Al foldon trimerization domain based on phage T4 fibritin followed by a c-myc epitope and a hexa-repeat histidine tag were introduced to the C-termini of both S-D614 and S-D614G as described previously[2].

The plasmids of S-D614 and S-D614G were transiently transfected into HEK293 Freestyle cells with polyethylenimine (PEI, linear, 25 kDa, Polysciences, U. S. A.) at a ratio of DNA: PEI = 1:2. The transfected cells were incubated at 37 ºC with 8 % $CO_2$ for six days. After pelleting cells by centrifugation at 4000 rpm for 30 min, the medium was harvested and filtered through a 0.22-µm cutoff membrane (Satorius, France). The cleared medium was incubated with HisPur Cobalt Resin (Thermo Fisher Scientific, U. S. A.) in binding buffer (50 mM Tris-HCl (pH 7.6), 300 mM NaCl, 5 mM imidazole and 0.02% $NaN_3$) at 4ºC overnight. The resin was iteratively washed with wash buffer (50 mM Tris-HCl (pH 7.6), 300 mM NaCl, 10 mM imidazole) and the target protein was subsequently eluted by elution buffer (50 mM Tris-HCl (pH 7.6), 150 mM NaCl, 150 mM imidazole). The protein was concentrated and loaded into a size exclusion chromatography (SEC) column (Superose 6 increase 10/300 GL; GE Healthcare, U. S. A.) with a running buffer containing 50 mM Tris-HCl (pH 7.6), 150 mM NaCl, 0.02 % $NaN_3$ for further purification. The protein concentrations were determined by using the UV absorbance at 280 nm using an UV-Vis spectrometer (Nano-photometer N60, IMPLEN, Germany).

#### Cryo-EM sample preparation of S-D614G and data collection

Three microliters of purified protein were applied onto 300-mesh Quantifoil R1.2/1.3 holyed carbon grids. The grids were glow-charged at 20 mA for 30 s. After 30-s incubation, the grids were blotted for 2.5 s under 4 ºC with 100% humidity, and vitrified using a Vitrobot Mark IV (ThermoFisher Scientific, U. S. A.).

Data acquisition was performed on a 300 keV Titan Krios microscope equipped with a Gatan K3 direct detector (Gatan, U. S. A.) in a super-resolution mode using the EPU software (ThermoFisher Scientific, U. S. A.). The cryo-EM data were collected in a movie mode with a defocus range of -0.8 to -2.6 μm at a magnification of 81000 x, which corresponded to a pixel size of 0.55 Å. A total dose of 48 $e^-/Å^2$ was distributed over 50 frames with an exposure time of 1.8 second. The dataset was collected with an energy filter (slit width: 15-20 eV), and the dose rate was adjusted to 8 $e^-$/pix/s.

Image processing and 3D reconstruction

All 2x binned super-resolution movies were processed by Relion-3.0[3] with dose-weighting and 5x5 patch-based alignment using the GPU-based software MOTIONCOR2[4]. All motion-corrected micrographs were further processed by cryoSPARC v2.14[5]. Contrast transfer function (CTF) estimation was performed by patch-based CTF. The micrographs that were considered by the script "CTF_fit_to_Res" as satisfactory (between 2.5 and 4 Å) were used for particle picking. A small subset of micrographs was used for unbiased template-free blob picker within cryoSPARC. The picked particles were extracted with a box size of 192 pixels (2x2-binned), followed by iterative 2D classifications with visual inspections to remove junk particles.

To identify conformational heterogeneity of S-D614G trimer, ~660k particles were initially classified by *ab-initio* reconstruction with a C1 symmetry (class = 5), followed by heterogeneous refinement to generate five distinct classes (class = 5). One of the 3D classes showed a single RBD-up conformation (388,400 particles), whereas the remaining classes showed poorly defined cryo-EM maps, which were therefore excluded from the subsequent data process. The particles that corresponded to the single RBD-up conformation were used for further processing by using non-uniform 3D refinement (NU-refinement) with a C1 symmetry. This yielded a 3.1 Å cryp-EM map according to the Fourier shell correlation (FSC) = 0.143. The map and mask from the NU-refinement were used for the 3D variability analysis (3DVA) as part of cryoSPARC, which generated five clusters for further heterogeneous refinement (class = 5). The particle images from each 3D class were unbinned and re-extracted with a box size of 384 pixels. The full-resolution particle stacks were used for the NU-refinement, local CTF refinement, a second round of NU-refinement to generate five final cryo-EM maps, including three “one RBD-up” conformations (3.5, 3.6 and 3.6 Å) and two “two RBD-up” conformations (3.5 and 3.4 Å). The local resolution analysis for was calculated using ResMap[6].

Model building and refinement

An initial model was generated by Swiss-Model[7] using the PDB structure 6XM3 as a template. The atomic coordinates were divided into individual domains and manually fit into the cryoEM maps of S-D614G by using UCSF-Chimera[8], UCSF-ChimeraX[9] and Coot[10]. After iterative refinements, the structural models were refined by the real-space refinement module within Phenix[11]. 22 previously reported N-glycosylation sites were examined further to identify additional cryo-EM densities protruding from the corresponding asparagine side-chain, which implied the presence of N-glycan moieties. In cases where additional EM densities were clearly visible, atomic models of N-linked glycans were built onto the asparagine side-chains by using the module "Glyco" within Coot[10]. The final model was assessed by MolProbity[12]. Structural visualization and rendering of structural representations were achieved by using a combination of UCSF-Chimera, UCSF-ChimeraX and Pymol (Schrodinger Inc. U. S. A.).

Glycopeptide analysis

The glycopeptide analyses of S-D614 and S-D614G were carried out using the protocol as described previously[2] with the following modifications. First, the protein was digested by trypsin and chymotrypsin (Promega) with a protein to protease ratio of 25:1. Second, the digests were separated using a segmented gradient of 5 to 40% solvent B over 200 minutes. The MS/MS data were identified by the Byonic software (v.3.9.6). The ion chromatograms of accepted peptide match precursors were extracted at 5 ppm to record the peak area of each unique glycopeptide and used to compare the relative amount of glycoforms in two samples.

Negative staining electron microscopy analysis (NSEM)

Four microliters of S-D614 and S-D614G that were treated at different temperatures and durations – fresh (day 0), fresh samples incubated at 50 ºC/60 ºC for 30 min, and fresh samples incubated at 4 ºC/37 ºC for six days –were used to prepare negative staining EM grids at a concentration of 50 μg/mL. The carbon-coated grids were glow-discharged with 25 mA for 30 s. After staining with 0.2% uranyl formate (UF), the grids were blotted and dried at the air for one day. Images were collected by using a FEI Tecnai G2-F20 electron microscope at 200 keV (FEI, the Netherlands). A magnification of 50000x was used, corresponding to a pixel size of 1.732 Å. All datasets were processed by cryoSPARC v2.14, including patch-CTF estimation, particle picking/extraction, 2D classification and ab-initio 3D reconstruction. The number of "intact" spike particles in each micrograph (defined by the number of particles used in 3D reconstruction) was extracted by the function "Manually curate exposures" within cryoSPARC. The numbers were exported to GraphPad Prism 8 (GraphPad, U. S. A.) for further statistical analyses. The 3D models of individual experimental conditions were visualized by using UCSF-ChimeraX.

Differential scanning calorimetry (DSC)

The thermal unfolding of S-D614 and S-D614G were analyzed by DSC (Automatic MicroCal PEAQ-DSC. Malvern, United Kingdom). The protein concentrations were set to 0.2 mg/mL in 50 mM Tris-HCl (pH 7.6), 150 mM NaCl, and 0.02 % $NaN_3$. The temperature was ramped up from 10 to 90 ºC at a rate of 200 ºC/h. The resulting data were baseline-corrected by the built-in software of MicroCal PEAQ-DSC, and exported to GraphPad Prism 8 to generate graphs.

Differential scanning fluorimeter (DSF)

All experiments were performed in Tycho NT.6 (NanoTemper Technologies). Fresh samples, from biological replicates, were ten-times diluted in 100 mM sodium acetate (pH 4.0 and pH 5.0), MES (pH 6.0), HEPES (pH 7.0) or Tris (pH 7.6 and pH 8.6) supplement with 150 mM NaCl at the final concentration of 0.15 mg/mL. The temperature range was set to 35-95ºC and the scanning rate was 30ºC/min. All data were processed in the internal software from NanoTemper, and the statistics of experimental parameters analyzed by GraphPad Prism 8.

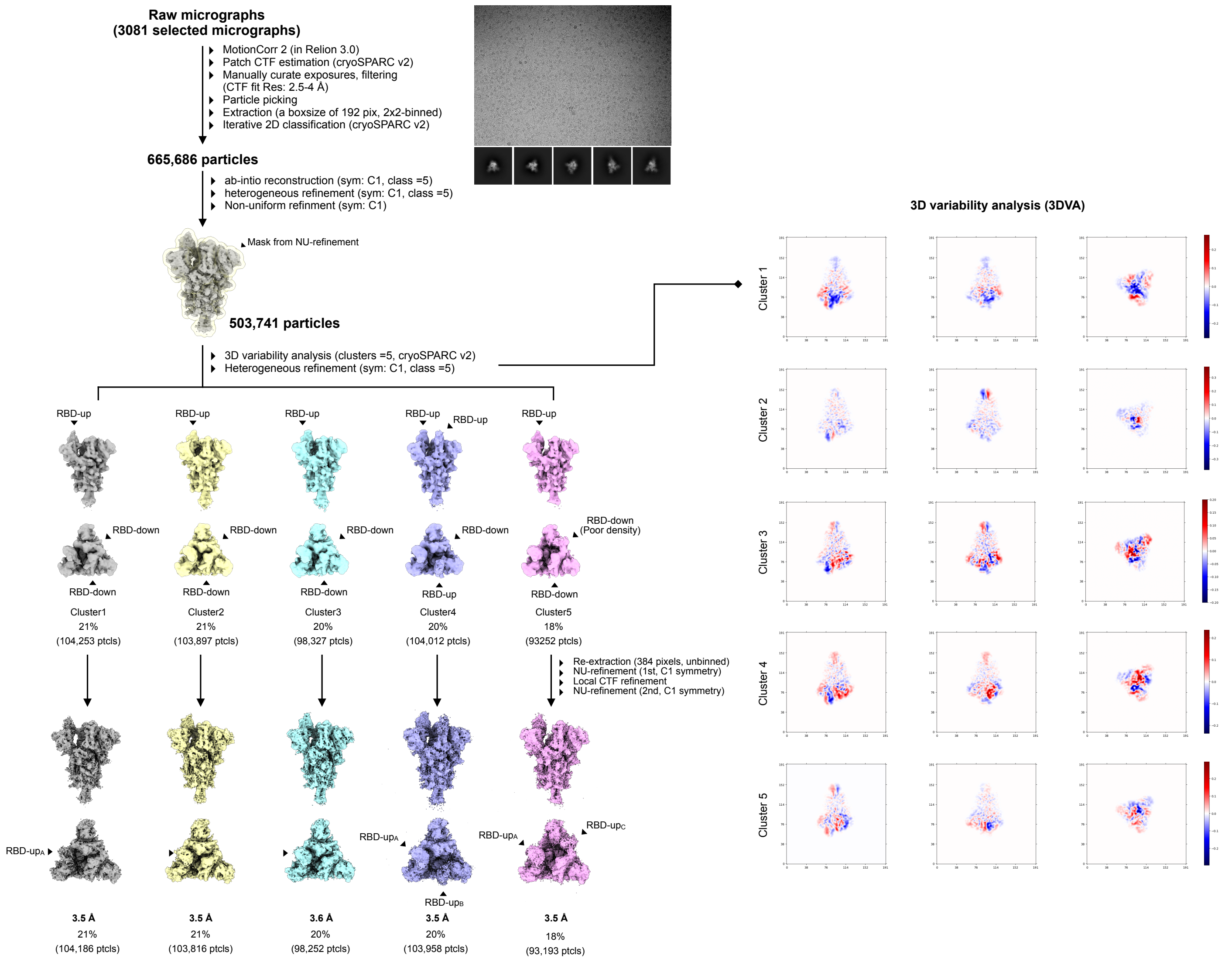


**Fig. S1. Workflow for cryo-EM data processing.**

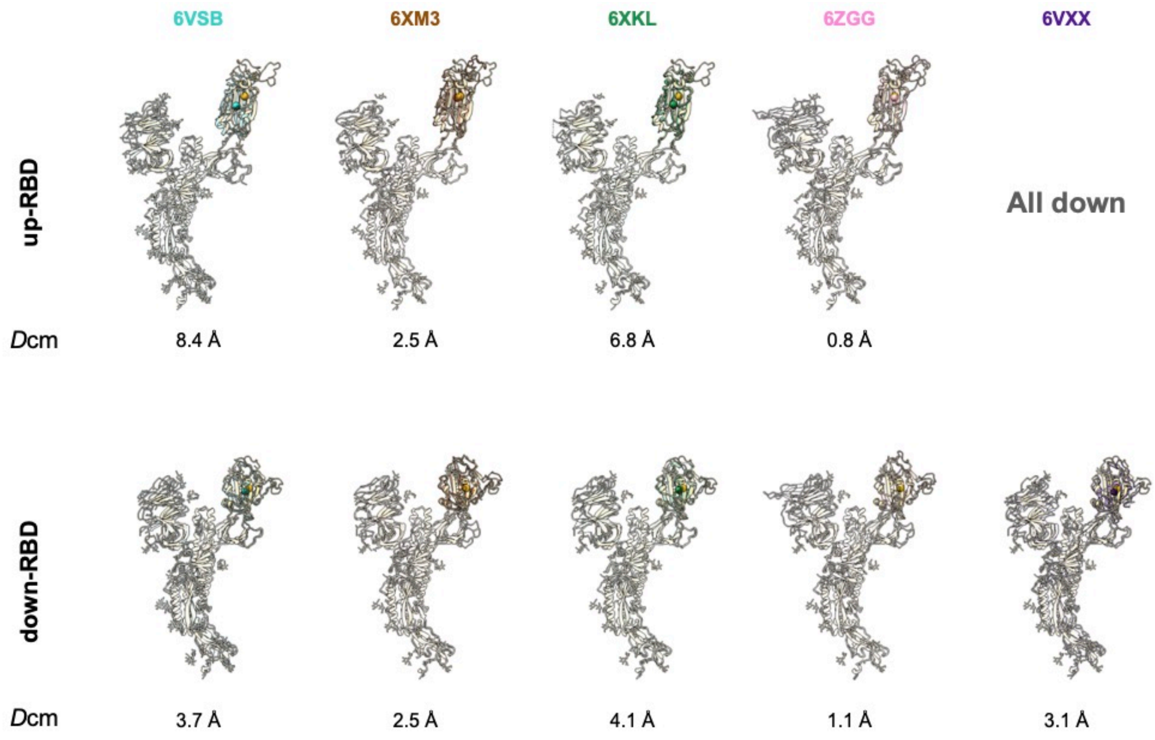


**Fig. S2. Comparison of RBD orientations of reported S-D614G and S-D614 structures.** Cartoon representations of RBD-up and RBD-down protomers derived from previously reported cryo-EM structures and our current study. The PDB accession codes of individual structures are shown above. The PDB entries of the representative spike variants are 6VSB (S-D614 in an one RBD-up), 6VXX (all RBD-down, closed state), 6XM3 (one RBD-up, at pH 5.5), 6XKL (engineering thermal-stable mutant, HexaPro, one RBD-up), and 6ZGG (furin-cleaved, one RBD-up). All protomers were superimposed with respect to the S2 domains of S-D614G of this study. After the alignment with respect to the S2 domain, the displacement between the centers of mass (Dcm; indicated by spheres) of previously reported RBD structure and ours is calculated by UCSF-Chimera, which is indicated below each structure.

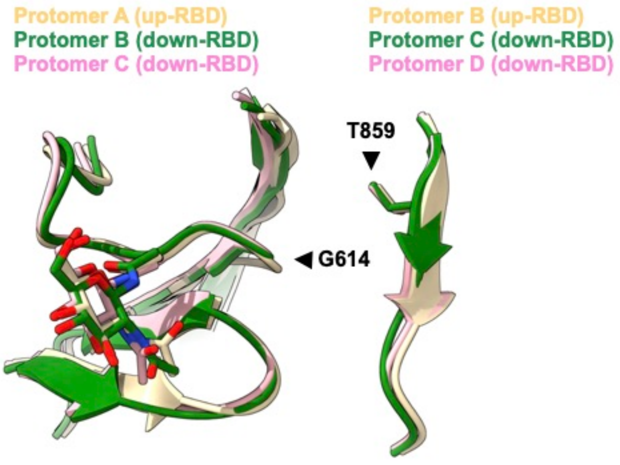


**Fig. S3. Superposition of the structural elements surrounding the D614G mutation site from within the three protomers (chains) of S-D614G (chains A, B, and C).** The results indicated that local structures of the mutation site are essentially identical, despite the difference of their corresponding RBD orientations. The protomers are colored in wheat (protomer A, RBD-up), green (protomer B, RBD-down), and pink (protomer C, RBD-down), respectively. The first GlcNAc moiety that is linked to the side chain of N616 is shown in sticks.

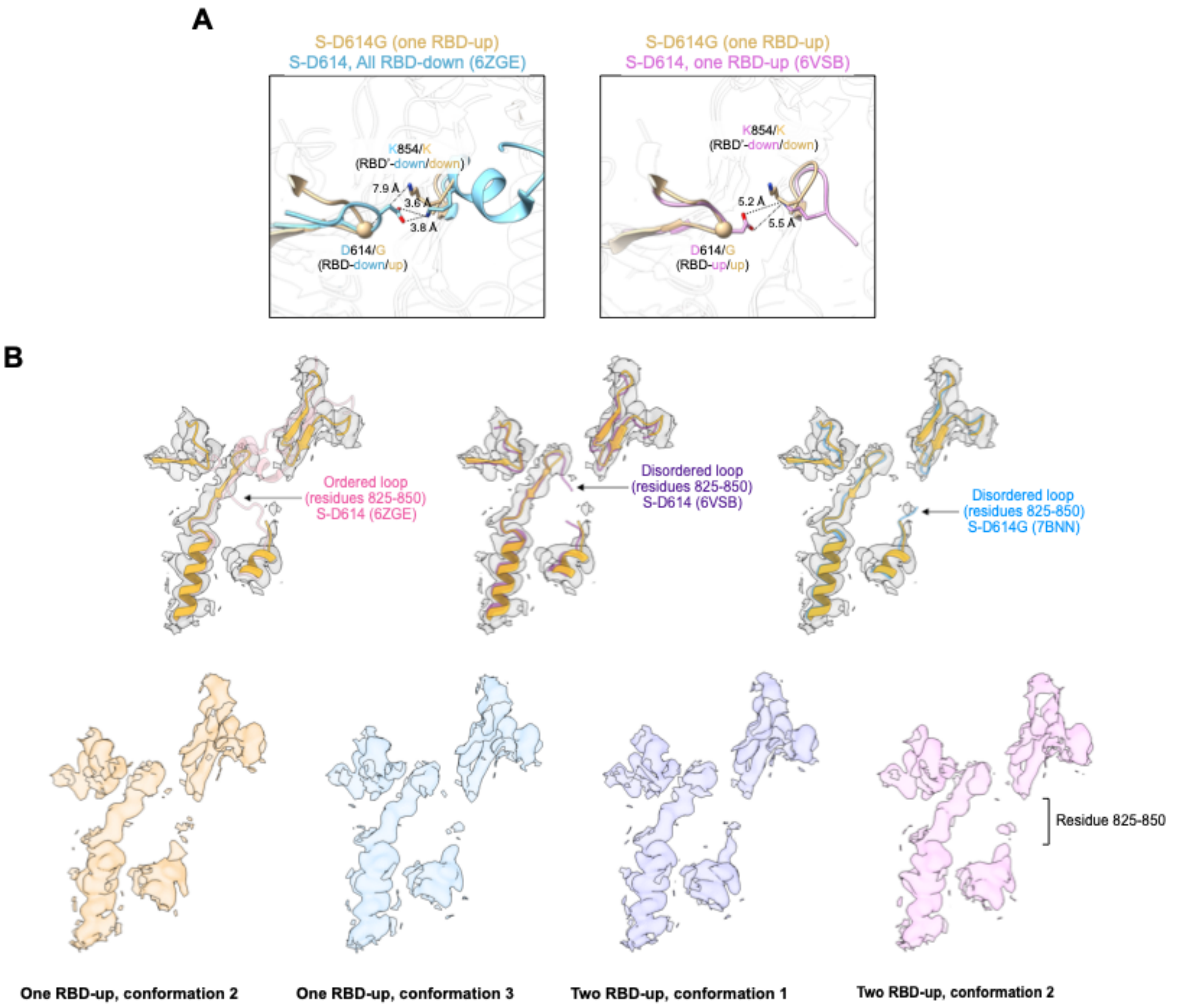


**Fig. S4. Conformational disorder around the D614G mutation site.** (A) Structural comparison of the inter-protomer interaction between S-D614G and S-D614 (two conformations: all RBD-down and one RBD-up). The dash lines indicate the distance between G614 (or D614) and K854. The structure of S-D614G is colored in tan. The structures of S-D614 with all RBD-down (PDB code: 6ZGE; left panel) and with one RBD-up (PDB code: 6VSB; right panel) are colored in blue and pink, respectively. (B) The loop on which K584 resides is disordered in all the selected cryo-EM structures, which is manifested in the lack of defined cryo-EM maps for residues 825-850 as indicated on the right.

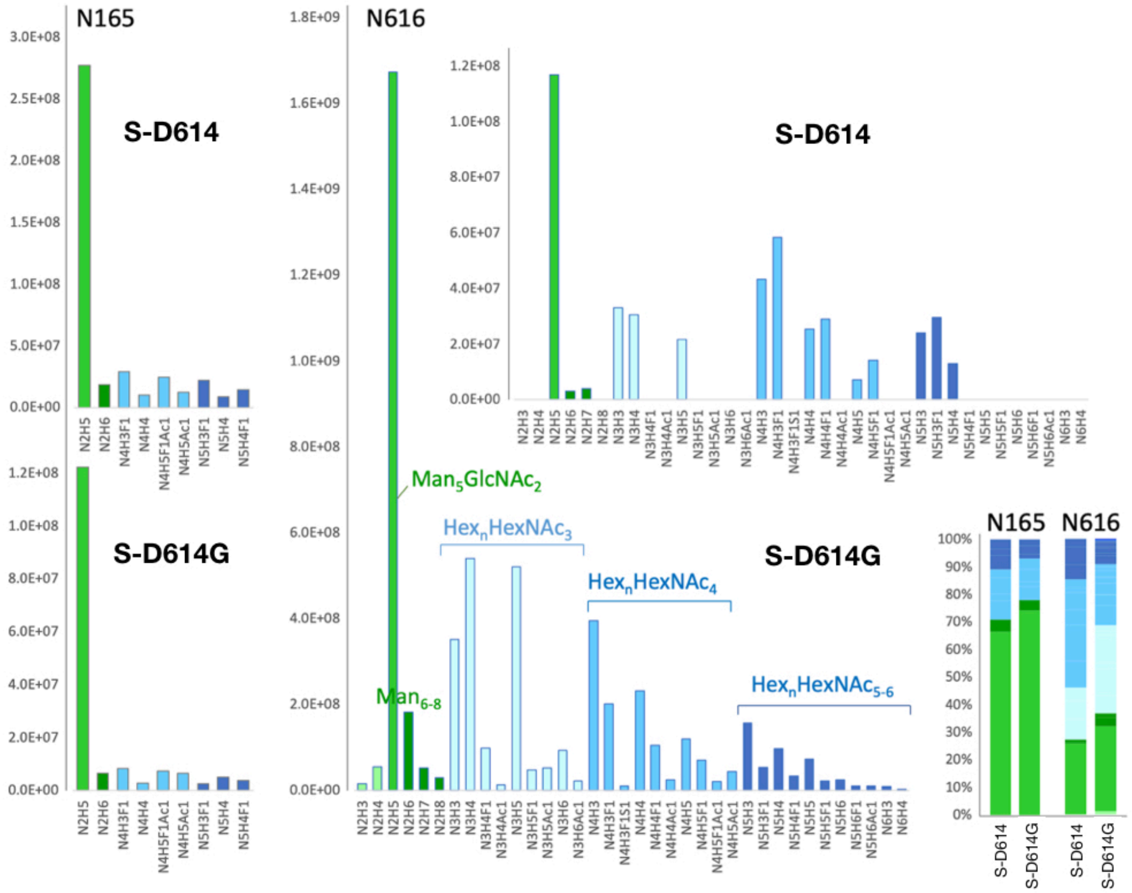


**Fig. S5. Glycoform distribution for N-glycosylation sites within the vicinity of D614G.** LC-MS/MS analysis of glycopeptides from digested S-D614 and S-D614G demonstrated an overall very similar site-specific glycosylation pattern consistent with previous reports by others. The glycosylation pattern for each of the three N-glycosylation sites located close to the mutated D614G site in 3D structures, namely N165, N234 and N616, remained largely unchanged compared to S-D614. However, the amount of the predominant $Man_5GlcNAc_2$ glycoform relative to other hybrid ($Hex_nHexNAc_{3,4}$) and complex type ($Hex_nHexNAc_{4\text{-}6}$) structures did increase slightly. It was further noted that the although sample preparation and analyses were performed similarly in parallel, the recovery of N616-glycopeptides from S-D614G was about an order higher, probably due to the amino acid substitution, whereas those of N165-glycopeptides were similar.

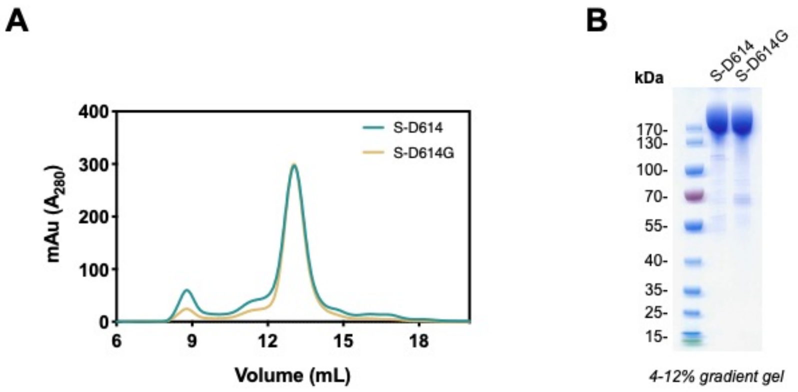


**Fig. S6. Purification of recombinant S-D614 and S-D614G.** (A) Overlaid SEC chromatograms of S-D614 (grass green) and S-D614G (yellow). (B) Coomassie Brilliant Blue stained SDS-PAGE, which illustrated the purity and integrity of the purified recombinant S-D614 and S-D614G.

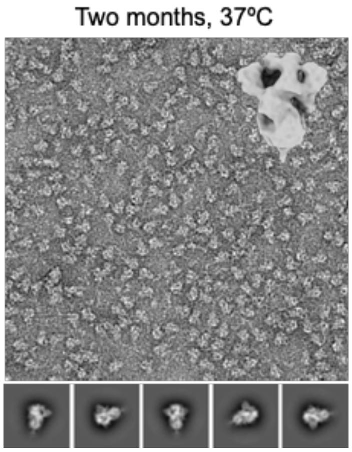


**Fig. S7. Representative NSEM image and 2D classes of S-D614G incubated at 37ºC for 60 days.** The 3D model derived from the NSEM data was shown on the upper right corner.

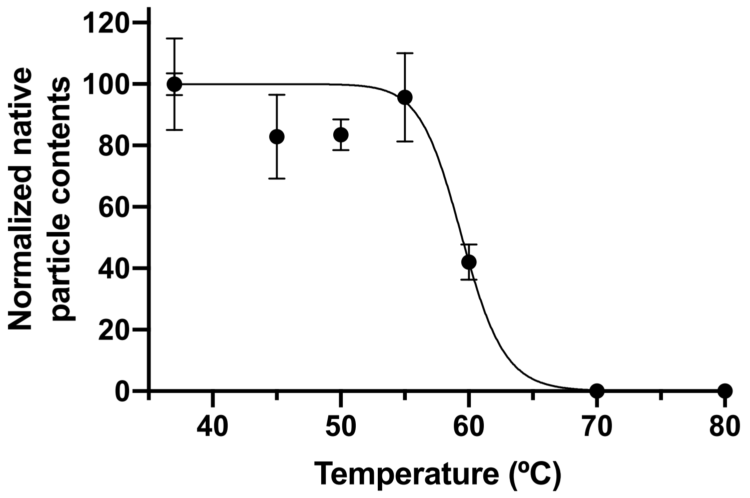


**Figure S8. Relative native-like particle number of S-D614G as a function of temperature.** All samples were incubated at the specified temperatures as indicated along the X-axis for 30 min prior to NESM grid preparation. The numbers of native-like particles in individual micrographs were normalized with respect to the average value of freshly prepared sample (37 ºC, day 0). The error bars correspond to the standard deviations of the numbers within the micrographs collected under the same conditions. For each temperature, between 50 and 60 NSEM micrographs were collected.

**Table S1. Parameters of cryo-EM data collection, processing and model validation.**

| | One RBD-up conformation 1 | One RBD-up conformation 2 | One RBD-up conformation 3 | Two RBD-up conformation 1 | Two RBD-up conformation 2 |
|---|---|---|---|---|---|
| **Data collection and processing** | | | | | |
| Microscope | | | Titan Krios (Gatan K3 Summit camera) | | |
| Voltage (keV) | | | 300 | | |
| Mode | | | Counting | | |
| Magnification | | | 81,000x | | |
| Total dose ($e^-/Å^2$) | | | 48 | | |
| Defocus range (μm) | | | 0.8-2.6 | | |
| Pixel size (Å) | | | 1.1 (2x binned) | | |
| Final used particles | 104,186 | 103,816 | 98,252 | 103,958 | 93,193 |
| Symmetry | C1 | C1 | C1 | C1 | C1 |
| Map Resolution (Å) | 3.5 | 3.6 | 3.6 | 3.5 | 3.4 |
| **Model composition** | | | | | |
| Non-hydrogen atoms | 24,959 | 24,949 | 24,847 | 24,835 | 24,905 |
| Protein residues | 3,051 | 3,055 | 3,041 | 3,047 | 3,058 |
| Ligands | 82 | 80 | 79 | 75 | 74 |
| MolProbity score | 1.81 | 1.79 | 1.85 | 1.74 | 1.79 |
| *Ramachandran (%)* | | | | | |
| Favored | 94.67 | 94.08 | 93.61 | 95.20 | 94.42 |
| Allowed | 5.29 | 5.88 | 6.32 | 4.77 | 5.44 |
| Outliners | 0.03 | 0.03 | 0.07 | 0.03 | 0.13 |
| Rotamer outliners (%) | 0.00 | 0.00 | 0.04 | 0.00 | 0.00 |
| Clashscore | 8.21 | 7.12 | 7.90 | 7.35 | 7.52 |
| *r.m.s. deviations* | | | | | |
| Bond length (Å) | 0.003 | 0.003 | 0.004 | 0.003 | 0.003 |
| Bond angles (º) | 0.600 | 0.586 | 0.593 | 0.557 | 0.573 |
| **Data Deposition** | | | | | |
| PDB code | 7EAZ | 7EB0 | 7EB3 | 7EB4 | 7EB5 |
| EMDB code | 31047 | 31048 | 31050 | 31051 | 31052 |

**Table S2. Parameters of Negative stain electron microscopy (NSEM) data collection and processing.**

| | S-D614 | | | | | S-D614G | | | | | |
|---|---|---|---|---|---|---|---|---|---|---|---|
| | Day 0 | Day 0<br>50ºC<br>30 min | Day 0<br>60ºC<br>30 min | Day 6<br>37ºC | Day 6<br>4ºC | Day 0 | Day 0<br>50ºC<br>30 min | Day 0<br>60ºC<br>30 min | Day 6<br>37ºC | Day 6<br>4ºC | Day 60<br>37ºC |
| **Data collection** | | | | | | | | | | | |
| Microscope | Tecnai G2-F20 | | | | | | | | | | |
| Voltage (keV) | 200 | | | | | | | | | | |
| Magnification | 50,000 x | | | | | | | | | | |
| Total dose ($e^-$/Å$^2$) | 30 | | | | | | | | | | |
| Micrographs collected | 54 | 54 | 61 | 56 | 57 | 58 | 64 | 55 | 56 | 55 | 55 |
| Pixel size (Å) | 1.732 | | | | | | | | | | |
| Initial number of particles | 65,897 | 97,728 | 111,929 | 96,971 | 104,714 | 103,866 | 109,150 | 98,131 | 98,540 | 93,279 | 92,277 |
| Number of particles used for EM map building | 39,122 | 25,688 | 3,985 | 35,869 | 1,894 | 52,784 | 48,639 | 21,039 | 50,625 | 44,106 | 52,181 |
| Symmetry | C1 | | | | | | | | | | |

**Table S3**. **Summary of DSC analysis of S-D614 and S-D614G after different treatments**

| Condition | Sample | Enthalpy of unfolding (kcal mol$^{-1}$) | $T_{m1}$ (°C) | $T_{m2}$ (°C) |
|---|---|---|---|---|
| **Fresh** | S-D614 | 1580 | | 66.9 |
| | S-D614G | 2250 | | 68.8 |
| **37 °C, 6 days** | S-D614 | 997 | | 67.0 |
| | S-D614G | 1560 | | 68/9 |
| **4 °C, 6 days** | S-D614 | 853 | 48.2 | 66.8 |
| | S-D614G | 2120 | 49.0 | 68.9 |
